# Genome-wide screen uncovers novel host factors for L-A virus maintenance and a mutualistic-symbiosis relationship in yeast

**DOI:** 10.1101/2025.05.20.655038

**Authors:** Wan-Yi Hsiao, Chung-Shu Yeh, Hsin-I Liu, Luh Tung, Tien-Hsien Chang

**Author notes:** Corresponding Author: Tien-Hsien Chang, Genomics Research Center, Academia Sinica, 128 Academia Road, Section 2, Nankang, Taipei, 115, Taiwan.

## Abstract

Viruses are often regarded as obligate intracellular parasites that exploit host resources for their own propagation. However, emerging evidence suggests that virus–host interactions can be more complex than simple antagonism. Here, we performed a genome-wide screen in *Saccharomyces cerevisiae* to identify host factors required for the maintenance of the L-A double-stranded RNA virus, a persistent and non-lytic resident of most laboratory yeast strains. Using two complementary mutant collections encompassing ∼6,000 yeast genes (∼93% genome coverage), we identified 96 host genes essential for L-A maintenance, spanning diverse biological functions. Transcriptome profiling revealed that the presence of L-A virus alters the host stress-response gene expression program. Strikingly, competitive fitness assays under environmental stress conditions showed that L-A enhances host stress tolerance, revealing a previously unrecognized mutualistic relationship. Together, our findings redefine the L-A–yeast interaction as a form of stable mutualism and highlight the utility of functional genomics and systems-level approaches in uncovering hidden dimensions of virus–host coevolution.

**Graphical Abstract:** 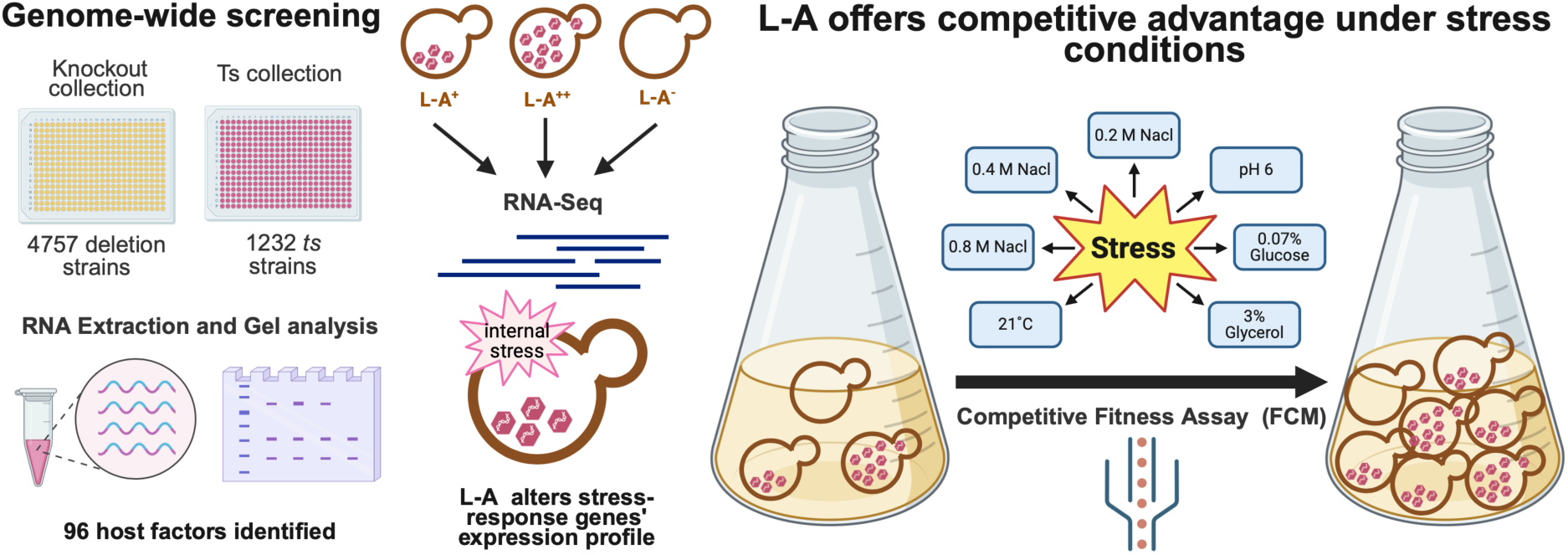

## Introduction

The interaction between virus and its host is an unending battle, in which virus often quickly evolves strategies to combat the host defense system. Yet, relative to their host genomes, virus genomes are considerably smaller, thus during their replication in the cells, viruses must recruit host factors to complete their replication machinery, only parts of which are encoded in the viral genomes. Conversely, the host would develop various ways to limit the virus in the cell (Koonin 2017). How and why some viruses successfully achieve “permanent residency” in their hosts is an important biological and evolutionary question highly relevant to human diseases.

Several virus-like elements exist in the budding yeast *Saccharomyces cerevisiae*, including the well-studied intracellular double-stranded RNA (dsRNA) viruses L-A and its satellite M1 (Wickner 1996). Practically all laboratory yeast strains carry the L-A virus (Wickner 1996), which has a single 4.6-kb dsRNA genome consisting of two overlapping open reading frames (ORFs). The first ORF encodes Gag, the major coat protein of the L-A virus. The second encodes the Gag-Pol fusion protein, an RNA-dependent RNA polymerase, whose complete coding region is established by a -1 ribosomal frameshift about 130 nucleotides upstream of the Gag-ORF’s stop codon. In contrast, the M1 virus has a 1.8-kb genome encoding only a 32-kd toxin/immunity precursor protein, which is subsequently processed and secreted as mature toxin (Wickner 1993). Together, L-A and M1 viruses form one of the yeast “killer” systems in which the M1 virus, whose replication is dependent on L-A, produces toxins fatal to cells not harboring the M1 virus (Wickner 1986). It is this distinct assayable phenotype that makes these dsRNA viruses amenable to genetic analysis and therefore provides a good model system for studying virus-host interaction in eukaryotes (Pagé et al. 2003; Zhao 2017; Pieczynska et al. 2017).

The L-A and the M1 viruses, whose genomes are encapsidated into virus-like particles (VLPs), appear to stably persist in the host’s cell cytoplasm without lysing the host cell or yielding detectable symptom (Wickner 1996). So far, there is no report of extracellular mode of transmission for the L-A and the M1 viruses and so it is generally believed that these viruses are spread through sexual mating or heterokaryon formation. Thus, historically, scientists took advantage of the toxin-producing property of the M1 virus to genetically identify host genes that are essential for maintenance, expression, or replication of the M1 dsRNA virus and, by extrapolation, the L-A virus too. For example, *MAK* genes are required for *<u>ma</u>*intenance of the *<u>k</u>*iller phenotype. In contrast, mutations in *SKI* genes yield the "*<u>s</u>*uper*<u>ki</u>*ller" phenotype, indicating that wild-type *SKI* genes function in controlling (or repressing) virus replication (Brown et al. 2000). Extensive characterization of these host factors over the years has yielded a wealth of knowledge regarding how these dsRNA viruses multiply in the cell.

It makes logical sense for the host to retain both the L-A and the M1 viruses at the same time in the face that the RNAi system is lost in yeasts of the *sensu stricto* clade, which includes *S. cerevisiae* (Drinnenberg et al. 2011). Presumably this is because the M1-encoded toxin more than offsets the disadvantage of losing RNAi (Drinnenberg et al. 2011). Yet, it remains baffling as to why many yeast strains contain only the L-A virus without its M1 satellite. As stated above, previous studies have already identified a number of host factors, suggesting a substantial diversion of host resources to the virus-propagation purpose. Thus, on the energy balance sheet, persistently keeping the L-A virus alone in the host appears to defy explanation. One argument would be that without the RNAi system, the host simply cannot efficiently clear the L-A virus. Here we entertain an alternative hypothesis that the L-A virus may have provided its host with heretofore unknown selective advantage, which would explain its residency in the host. To test this hypothesis, we performed an unbiased genome-wide screen solely on the basis of the L-A’s physical presence, aiming to identify host factors required for L-A’s maintenance. We identified a total of 96 host-factor genes, most are novel and were not found in the previous M1-based screens. The absence of these genes causes either a loss or elevated levels of the L-A virus. Transcriptomic and competitive fitness analyses further unveiled a mutualistic symbiosis between the L-A virus and its host under various stress conditions.

## Results and Discussion

### Identification of gene products functioning in L-A virus maintenance

To systematically identify gene products that function in L-A virus maintenance, we conducted a genome-wide screen of nearly all annotated genes in the budding yeast. A more recent study suggests a total of 6,275 protein-encoding genes exist in the budding yeast genome (Liu et al. 2017). Accordingly, we obtained two collections of *MAT***a** haploid yeast strains from C. Boone’s laboratory (University of Toronto). The first contains 4,742 nonessential-gene-deletion strains (Cherry et al. 2012) and the second 1,232 temperature-sensitive mutant strains, which correspond to 1,101 essential genes (Li et al. 2011). Together, these collections cover ∼93% of the annotated genes.

One major challenge to our goal is that L-A virus alone appears not to impart a scorable phenotype to the host, unlike the toxin-encoded M1 virus. To resolve this issue, we took a brute-force approach relying on, as a start, directly visualizing the L-A dsRNA via agarose-gel electrophoresis. We first inspected the presence of the L-A dsRNA in the collection of 4,742 nonessential-gene deletion strains, which were cultured in 96-deep-well plates to saturation and their total cellular RNAs were extracted for agarose gel electrophoresis (Fig. 1A and 1D). To impose rigor to this approach, we performed five rounds of elimination, each with an increasingly stringent visualization criterion. For example, in the first round, we found 613 strains with reduced levels of L-A dsRNA. These 613 strains were then taken again from the original collection for the second-round analysis, which cut the number down to 505. In the end, we identified 76 strains with apparently zero (Supplemental Fig. S1) and 42 strains with elevated levels of L-A dsRNA, respectively. We then performed PCR analysis to ascertain the bar-coded deletion in each strain. Finally, in each strain, we measured the abundance of the L-A dsRNA by RT-qPCR (Fig. 1B, C) and the presence or absence of the Gag protein by Western blotting analysis (Supplemental Fig. S2). This last step allowed us to eliminate practically all false positives. Ultimately, this exercise identified 45 genes, when deleted, resulted in a dramatic loss (>5 orders of magnitude) (Fig. 1B) and seven genes resulted in elevated levels (>1.5-fold increase) of the L-A dsRNA (Fig. 1C), respectively. The latter is in line with a report that *ski* mutations elevate the L-A level by 4- to 10-fold (Wickner 1992). Thus, the former group of 45 genes are involved in maintaining and the latter group (7) in suppressing the L-A virus, respectively, in the budding yeast (Supplemental Tables S1, S2).

**Figure 1.**
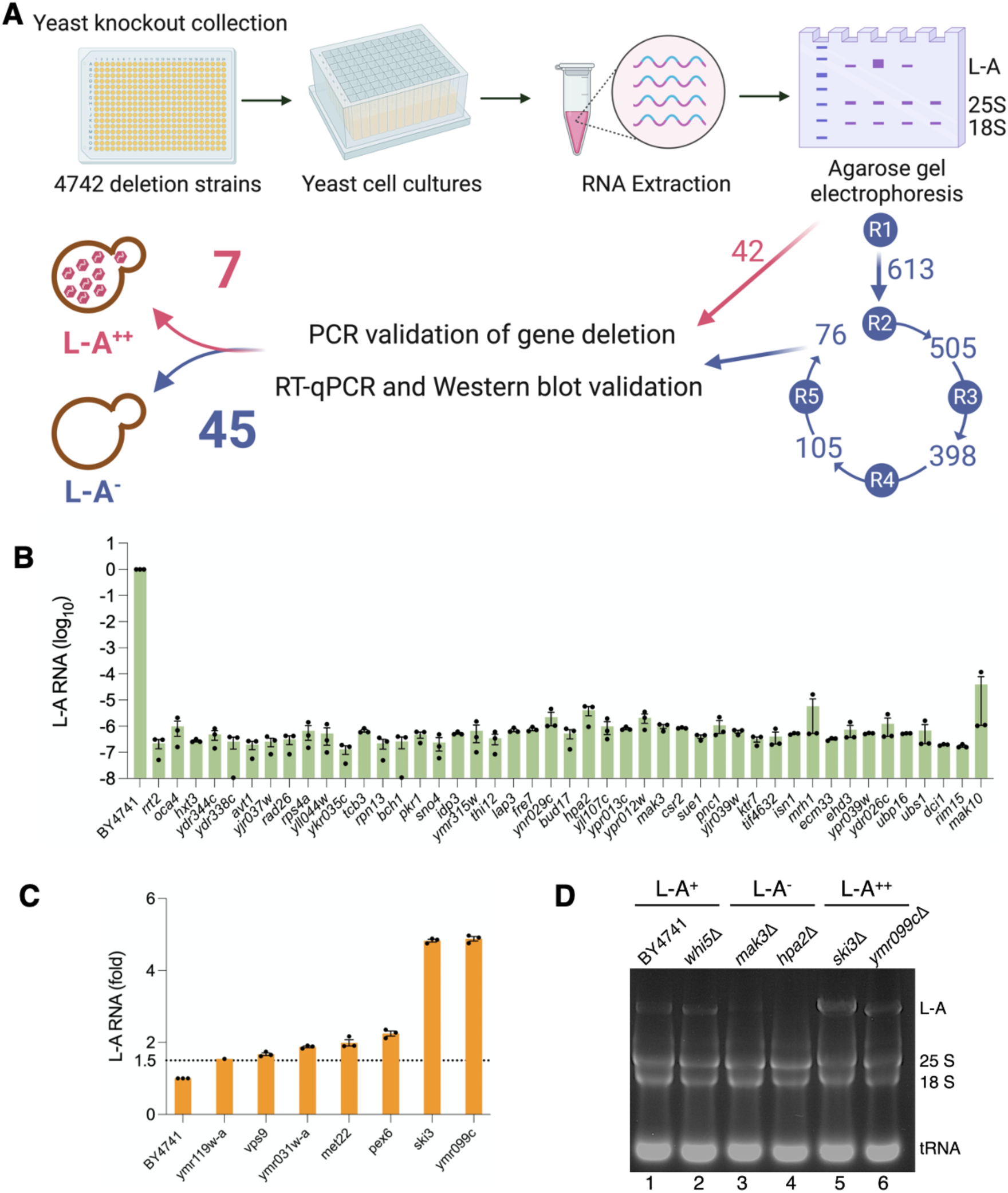
Genome-wide screen for L-A-maintaining gene products using a collection of the yeast-gene-knockout strains. (*A*) Schematic diagram for the genome-wide screen and validations. R1 to R5, five rounds of visual elimination; L-A^-^, strains with no detectable L-A dsRNA; L-A^++^, strains with elevated levels of L-A dsRNA. (*B*) RT-qPCR analysis of the 45 L-A^-^ finalists. BY4741 strain served as the wild-type control. (*C*) RT-qPCR analysis of the seven L-A^++^ finalists. *N* = 3 biological repeats, each with three technical repeats; error bars are ±SEM. (*D*) Representative agarose-gel-electrophoresis image showing wild-type level (lanes 1 and 2), absence of (lanes 3 and 4), and elevated levels of the L-A dsRNA (lanes 5 and 6).

It is of interest to note that several genes we uncovered have already been implicated in the life cycle of various RNA viruses, L-A virus included. For example, *MAK3* (Tercero et al. 1993) and *MAK10* (Lee and Wickner 1992) each of which encodes an acetyltransferase, are required for the L-A virus propagation (Tercero and Wickner 1992). In addition, we uncovered *HPA2*, which is predicted to encode an acetyltransferase as well. In contrast, *SKI3*, which encodes an exosome component, has an antiviral role in yeast (Hougan et al. 1989). Likewise, in a yeast-based brome mosaic virus (BMV) replication system, *OCA4* and *MET22* were found to play a promoting or a suppressing role in BMV RNA replication, respectively (Kushner et al. 2003). Furthermore, *DCI1*, which encodes a peroxisomal protein, affects tomato bushy stunt virus (TBSV) RNA recombination in a yeast model system (Serviene et al. 2005). Finally, *RAD26* is a homolog of the human ERCC6, which is up-regulated upon hepatitis C virus (HCV) infection (Wang et al. 2017). These cases thus suggest that most, if not all, of the uncovered genes are intimately tied to the L-A virus’s life cycle.

### Identification of essential gene products functioning in L-A virus maintenance

To identify essential genes for L-A maintenance, we first used several temperature-sensitive (ts) mutants to establish a general temperature-shift protocol (temperature, shift duration, and culture volume, etc.), seeking to strike a balance between rates of loss of the L-A virus and cell death. We then used this protocol (Fig. 2A) to screen a collection of 1,232 ts mutants via three rounds of visualization elimination (Fig. 2D) and RT-qPCR measurement of L-A dsRNA. In the end, we identified 24 mutant alleles that reduced L-A dsRNA level by >50% (Fig. 2B) and 20 mutant alleles that elevated the L-A dsRNA level (Fig. 2C). Among these genes, a number of them have been implicated in life cycles of various RNA and DNA viruses, e.g., BMV, TBSV, HCV, HIV, SARS-CoV-2, Epstein-Barr Virus (EBV), and Polyoma virus (Supplemental Tables S3, S4). Notably, we found that both *mob2-28* and *mob2-37* alleles reduced L-A dsRNA and that the same *mob2-28* allele was also reported to reduce TBSV replication in a yeast model system (Nawaz-ul-Rehman et al. 2013).

**Figure 2.**
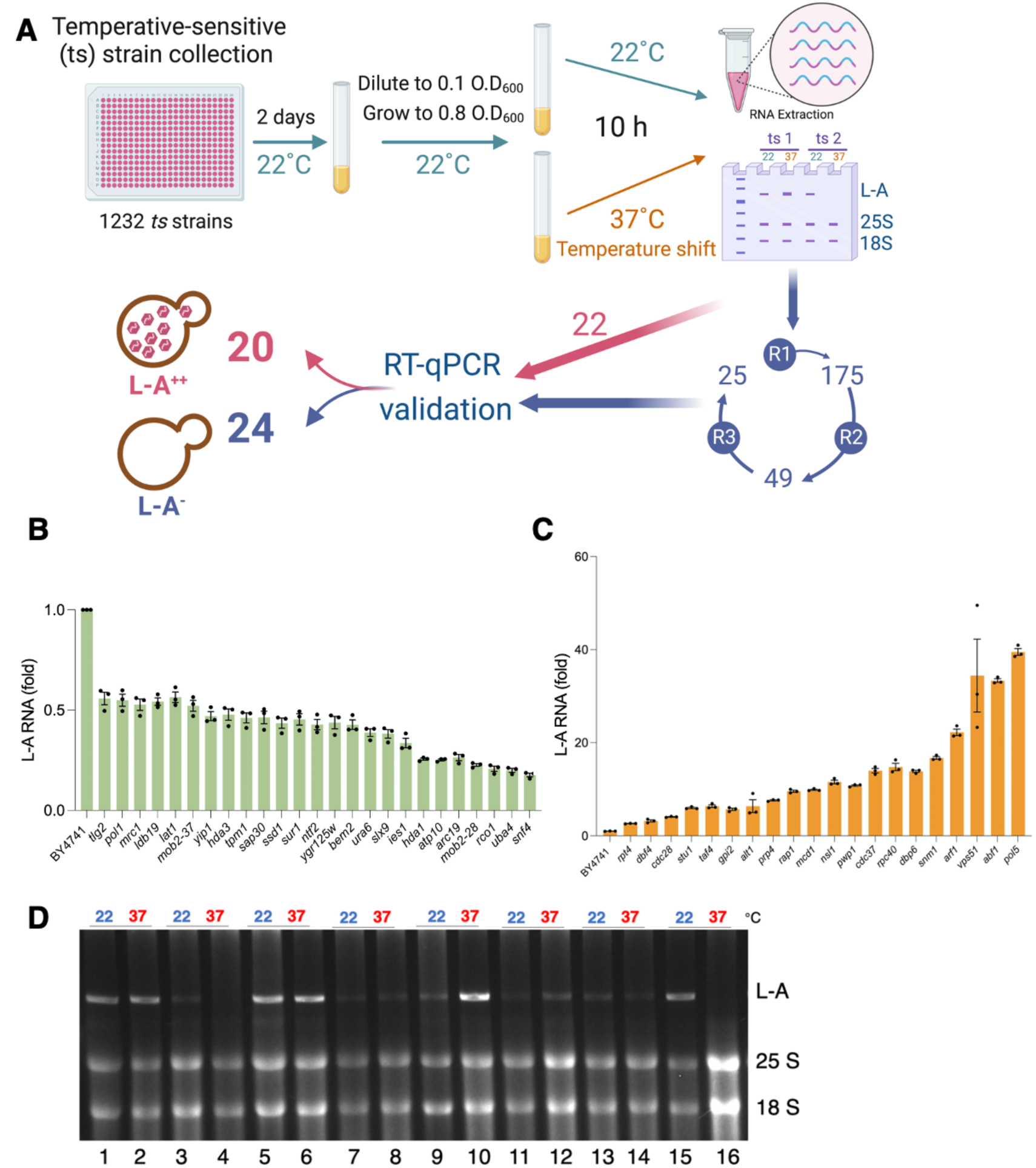
Genome-wide screen for L-A-maintaining gene products using a collection of temperature-sensitive mutants. (*A*) Schematic diagram for temperature-shift experiments and validations. Temperature shifted from 22°C to 37°C for 10 h. R1 to R3, three rounds of visual elimination; L-A^-^, strains with no detectable L-A dsRNA; L-A^++^, strains with elevated levels of L-A dsRNA. (*B*) RT-qPCR analysis of 25 ts mutant strains with reduced levels of the L-A dsRNA. The *MOB2* gene was independently identified twice through *mob2-37* and *mob2-28* mutants. BY4741 strain served as the wild-type control. (*C*) RT-qPCR analysis of the 20 ts mutant strains with reduced levels of the L-A dsRNA. BY4741 strain served as the wild-type control. *N* = 3 biological repeats, each with three technical repeats; error bars are ±SEM. (*D*) Representative agarose-gel-electrophoresis image showing data from eight ts mutant strains, each pair of analyses consists of total RNAs extracted from 22°C and 37°C cultures (e.g., lanes 1 and 2).

In summary, we identified a total of 96 genes whose deletion or mutant alleles impact on the L-A virus level. Among them, 69 are involved in maintenance and 27 in suppression of the L-A virus, respectively (Fig. 1A; Fig. 2A). Thus, at least 1.2% of the estimated 6,275 protein-encoding genes are involved in maintaining L-A virus in the budding yeast. This, however, is almost certainly an underestimate because we imposed a stringent criterion on screening the nonessential genes and relied only on a general protocol in screening the ts collection, which inevitably left out some candidates due to the heterogeneous nature of the ts phenotypes.

Gene ontology analysis (Fig. 3) revealed that the L-A-maintenance genes are enriched in small-molecule metabolic and catabolic processes, consistent with a view that metabolic control is critical to the host-pathogen interaction (Goodwin et al. 2015). In comparison, the L-A suppression genes are enriched in two notable categories of processes. The first category is related to cell-division-cycle processes, hinting that the L-A virus may interfere with the host’s cell-division-cycle (Subramani et al. 2018). The second category is related to nucleic-acid and RNA metabolic processes, in which the anti-viral Ski proteins are also classified. This may reflect a strategy of using exonucleases to suppress the L-A virus in the absence of the RNAi system in the budding yeast.

**Figure 3.**
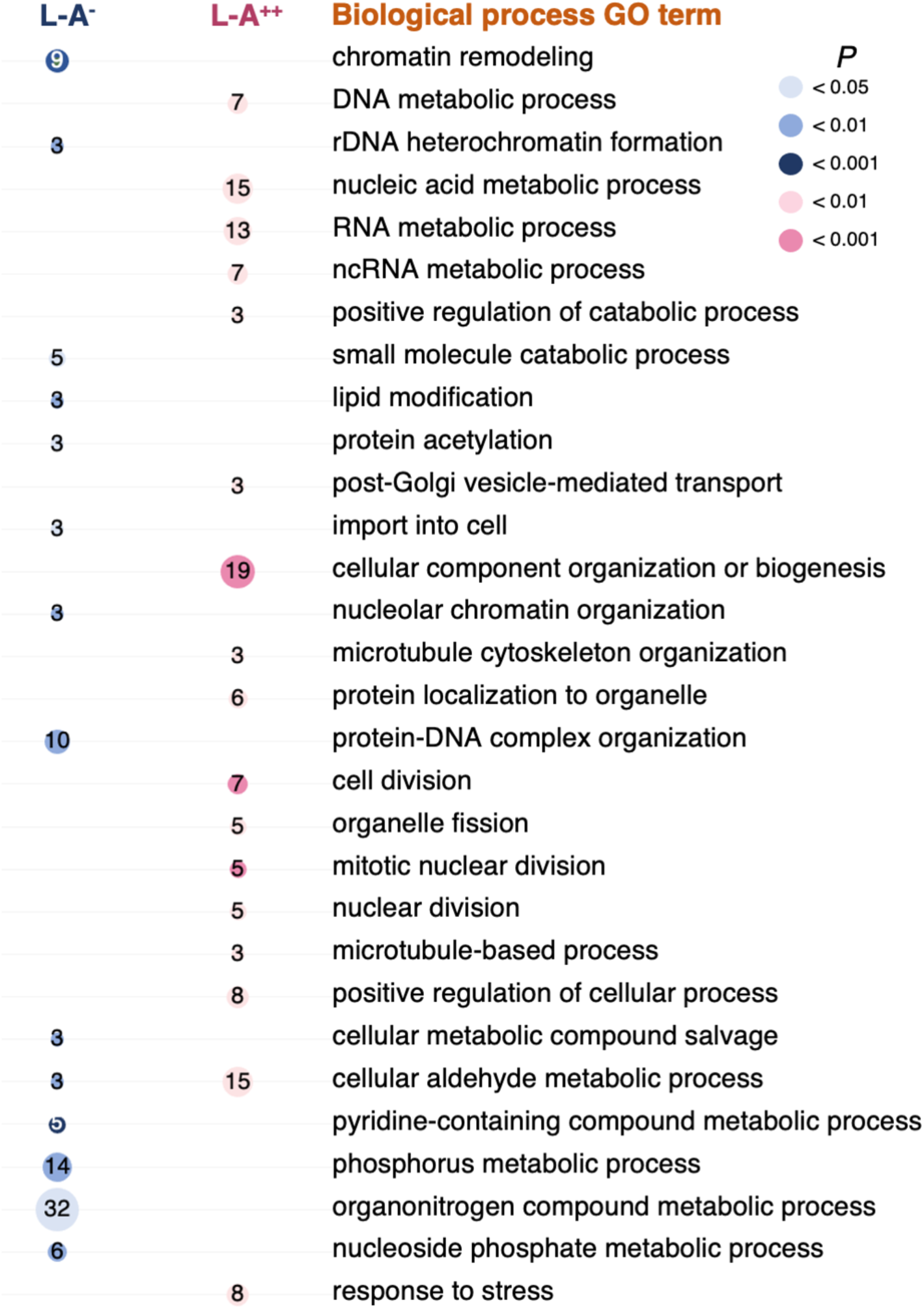
Gene ontology (GO) analysis of the identified genes in terms of biological processes. The size of the circle correlates with the number of genes classified to a specific biological process GO term. For example, in the L-A^++^ (elevated L-A dsRNA level) column, eight genes correspond to the “response to stress” GO term (bottom pink circle in the column). Statistical significance of the enrichment of each circle was computed to give a specific “colored” *P* value (upper right panel). Statistical analysis was done at LAGO (https://go.princeton.edu/cgi-bin/LAGO).

### L-A virus affects budding yeast’s gene-expression profile

To address L-A virus’s impact on the host physiology, we examined whether and how L-A’s abundance alters host’s gene-expression profile. In the collection of the nonessential-gene-deletion strains, there are 1,068 colonies of the wild-type BY4741 strain seeded as internal controls. During RT-qPCR analysis, we noticed that four (0.37%) colonies have no detectable and nine (0.84%) colonies have elevated levels of L-A dsRNA, respectively, perhaps due to sequential passages of the collection. Detailed transcriptomic analysis of the selected isolates showed that these isolates are virtually identical at the level of host’s genome (Materials and Methods; Table S7). Taking advantage of this observation, we compared transcriptomes among three BY4741 isolates that differ in their L-A virus loads, namely, the L-A^-^ (undetectable), the L-A^+^, and the L-A^++^ (3–5 fold higher than that of the L-A^+^) (Fig. 4A). These comparisons were done in a pairwise fashion, i.e., (1) L-A^-^ vs. L-A^+^; (2) L-A^-^ vs. L-A^++^; and (3) L-A^+^ vs. L-A^++^ (Fig. 4B). In Pair 1, we uncovered seven up-regulated and 22 down-regulated genes; in Pairs 2, nine up- and nine down-regulated genes; and in Pair 3, 11 up- and 4 down-regulated genes (Fig. 4B). Furthermore, three up-regulated and one down-regulated ones were shared in both Pair 1 and 2 comparisons; one up-regulated and one down-regulated genes in both Pair 2 and 3 comparisons (Fig. 4B). Notably, among all these genes, 10 up-regulated and 13 down-regulated ones are functionally related to environmental stress response (Gasch et al. 2000) (Supplementary Tables S5 and S6), suggesting that the presence of the L-A virus elicits a coordinated stress response in the host.

**Figure 4.**
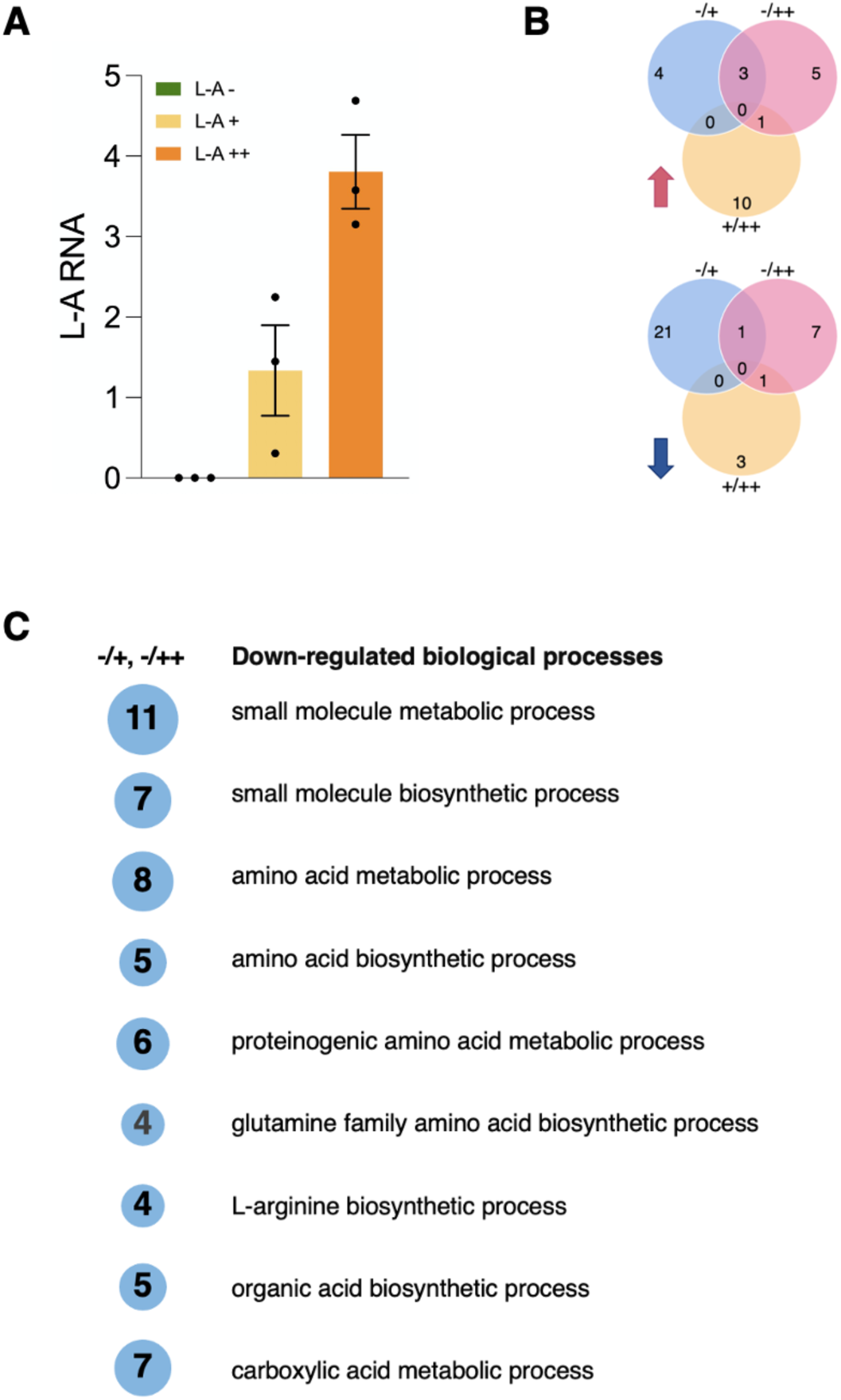
The L-A virus affects budding yeast’s gene-expression profile. (*A*) RT-qPCR quantification of the L-A^-^, L-A^+^, and L-A^++^ isolates. *N* = 3 independent cultures of each isolate; error bars are ±SEM. (*B*) Venn diagrams showing the number of up-regulated (top panel) or down-regulated (bottom panel) transcripts among the three isolates. (*C*) GO analysis in term of biological processes of the identified genes that are down-regulated in response to different L-A loads. The threshold of statistical significance was set at *P* < 0.001. The size of the circle corresponds to the number of genes within a specific biological process GO term. All analyses were done at was done at LAGO (https://go.princeton.edu/cgi-bin/LAGO).

Gene ontology analysis revealed that the L-A virus loads impact on a spectrum of genes involved in amino-acid metabolic and biosynthetic processes. These genes are enriched in the following GO terms: small molecule metabolic and biosynthetic process, amino acid metabolic and biosynthetic process, proteinogenic amino acid biosynthetic process, glutamine family amino acid biosynthetic process, L-arginine biosynthetic process, and carboxy acid metabolic process (Fig. 4C). On average, the change of host gene expression is about 4-fold (Supplementary Table S5 and S6). A previous study compared the transcriptomes of two strains, one harboring both L-A and M1 and the other without, also reported a small magnitude of gene-expression differences (McBride et al. 2013). Taken together, it seems that the coevolution between the L-A virus and its host may have been selected for a lesser impact on the host gene expression and may bear a close relationship to the general amino-acid metabolic processes (McBride et al. 2013).

### L-A virus offers competitive advantage to its host under stress conditions

Conceptually, the M satellite viruses provide a competitive edge to their host, because they encode secretable toxins (Wickner 1986). In contrast, the L-A virus alone offers no such benefit, yet it still requires nearly a hundred host genes (Fig. 1 and Fig. 2), a sizable fraction of the host genome, for its maintenance. So, why does the L-A virus not appear to exact a clear fitness cost to its host? This conundrum led us to entertain a hypothesis that L-A may tender a so-far unknown selective advantage to its host. Careful inspection of our transcriptomic analysis revealed that two-thirds of affected host genes are related to environmental stress response (Gasch et al. 2000). We therefore examined whether or not the L-A virus confers fitness benefit to its host under stresses.

To this end, we grew the L-A^-^, the L-A^+^, and the L-A^++^ strains under stresses and quantitatively measured their competitive fitness (see Materials and method). We chose eight conditions that mimic various physiological stresses, including salt concentration, pH value, glucose concentration, non-fermentable carbon source (glycerol), and temperature (Jerison et al. 2020). A trend emerged from this series of experiments is that the L-A harboring strains have higher fitness values, ranging from 1.4- to 3.5-fold increase, than that of the strain without L-A (Fig. 5). We ruled out the possibility that the observed fitness improvement is caused by the fluctuating L-A abundance over the course of experiment by RT-qPCR analysis (Supplemental Fig. S3). We noted that the L-A^++^ strain’s fitness level is lower than that of the L-A^+^ under the conditions of low glucose and the non-fermentable carbon source (Fig. 5).

**Figure 5.**
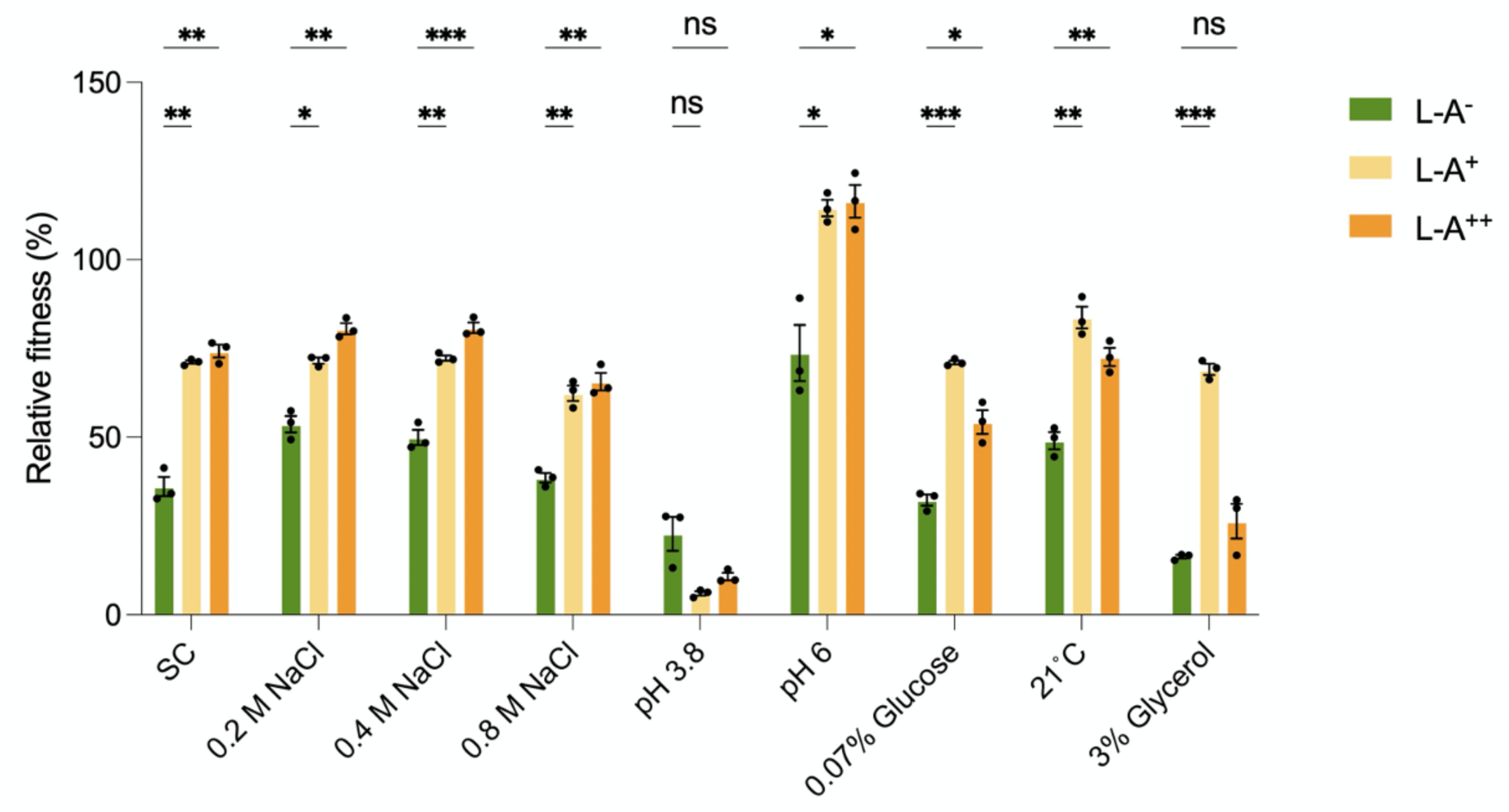
L-A virus offers competitive fitness advantage to its host under stress conditions. The flow-cytometry-based competitive fitness assay is detailed in Materials and methods and in the main text. Nine conditions were tested, including SC (synthetic complete medium). YPD rich medium cannot be used because it contains autofluorescent components. L-A^-^, green; L-A^+^, yellow; L-A^++^, orange. *N* = 3 biological repeats coupled with 3 technical repeats; error bars are ±SEM; \**P* < 0.05, \*\**P* < 0.01, \*\*\**P* < 0.001, and ns, no statistical significance

## Conclusion

In this study, we performed a genome-wide search for host factors required for the L-A virus maintenance in the budding yeast. Previous studies relied on the toxin-producing phenotype of the M satellite viruses, which inevitably would miss genes that are not functionally related to either M maintenance or toxin production. All told, we uncovered a total of 96 genes (Fig. 1B; 1C; 2B; 2C), most of them are novel, thus offering an opportunity to study these genes’ function in a system-biology fashion. Our transcriptomic analysis paints a picture that the L-A presence widely alters host’s stress-response gene expression, with 50 genes affected (Fig. 4B and Supplementary Table S5 and S6), supporting an adaptive co-evolution that may ultimately arrive at a virus-host symbiosis (McBride et al. 2013).

Conventionally, viruses are regarded as “selfish” intruders with a singular aim to propagate and expand themselves at the expense of the host. While some viruses remain dormant or even integrate themselves into the host genomes, their presence is still expected to exact a toll on the host. As such, the host invariably evolves a battery of antiviral strategies and *vice versa*, leading to an unending tug-of-war between the virus and its host. Here, the L-A virus represents an evolutionary enigma, in that, while diverting a wealth of cellular resources to achieve a permanent residency, its host seems not to have evolved or retained a systematic antiviral tool such as RNAi. Our finding that L-A improves the fitness of the host under stresses (Fig. 5) thus provides an explanation to the aforementioned puzzle.

Here, we offer a plausible model to account for the proposed mutualistic symbiosis relationship. The presence of a large number of L-A virus particles in the cytoplasm is expected to have already induced an internal stress response. Thus, upon external stress challenges, the host is able to mount a faster and fuller response. This is supported by two lines of evidence. First, eight out of a total of 27 gene deletions (Fig. 1A) and mutant alleles (Fig. 2A) that elevated the L-A levels (L-A^++^) are GO-classified in the stress-response processes (Fig. 3). This suggests that stress response is a part of the wider, but so-far hidden, web of anti-L-A defense system (Fig. 3) in the absence of the RNAi system. Indeed, taking out this defense system results in heightened levels of L-A (Fig. 1C; 2C). Second, transcriptomic analysis showed that the L-A presence affected the expression levels of environmental stress-response genes (Fig. 4B and Supplementary Table S5 and S6). Interestingly, we observed that an exceedingly high L-A load (L-A^++^) may ultimately overburden the host under two conditions, namely low glucose and non-fermentable carbon source (glycerol) (Fig. 5). This is reminiscent of a recent report that extremely high L-A load is lethal to the host grown in non-fermentable carbon source and at 37°C (Chau et al. 2023).

Broadly speaking, our finding is conceptually akin to a finding that chronic infection of mice with a γ-herpesvirus enhances resistance to *Listeria monocytogenes* and *Yersinia pestis* (Barton et al. 2007) and is, as well, compatible to the proposal that viruses may engage their hosts in conditional mutualism, mutualism, or symbiogenesis (Roossinck 2011; Roossinck and Bazán 2017). We now have a unique opportunity to investigate in detail the mechanism by taking advantage of yeast’s powerful genetic and biochemical tools.

## Materials and methods

### Screen for L-A virus maintenance genes

Non-essential-gene knockout (YKO) strains were cultured in liquid YPD (2% dextrose, 2% peptone and 1% yeast extract) medium in 96-deep-well plates for three days at 28°C. Standard hot phenol/chloroform procedure was used to extracted and isolated the total RNAs, which were run on agarose (0.8%) gels. We visualized the L-A dsRNA to identify the first-round candidates with apparently reduced or increased levels of L-A. These candidates were again pulled from the original source plates and re-analyzed. In total, we imposed five rounds of screening to rigorously eliminate false positives. To screen for essential genes, we obtained a collection of ts mutants from C. Boone’s lab (Li et al. 2011). These ts mutants are inviable at non-permissive temperatures (26–37°C), however their phenotypes are not uniform. For pilot experiments, we used four ts strains (*mak5*, *mak11*, *mak16*, and *mak21*) to arrive at a common temperature-shift protocol (Fig. 2A) for screening the entire collection. Briefly, each ts strain was grown to saturation at 22°C, from which equal number of cells (0.1 OD_600_) were seeded into two fresh cultures and grown to 0.8 OD_600_. One of the two cultures was then shifted to 37°C for 10 h, while the other remained at 22°C for the same duration of time. In the end, same number of cells were harvested from these two cultures for total RNA isolation and analyzed as described above.

### RT-qPCR analysis

RNA samples (5 µg) were converted into cDNA by Maxima First Strand cDNA Synthesis Kit (Thermo Scientific) following manufacturer’s instructions. Quantitative polymerase chain reaction (qPCR) was done by using StepOnePlus^TM^ Real-Time PCR System (7500 fast) with Fast SYBR^®^ Green Master Mix. Two oligonucleotide primers were used to amplify a part of Gag-coding region, i.e., LA1 (CCCGACAGTCTGCTTTAATGAAGG) and LA2 (ACCTGGGATACCGCAAGTGG). For YKO candidates, all L-A data were first normalized to that of the *ACT1* transcript, which served an internal control, and then normalized against the same *L-A/ACT1* value obtained from the wild-type BY4741 strain. For ts candidates, all L-A data first were normalized to that of the 18S rRNA, which remained constant under our temperature-shift protocol, and then normalized against the same *L-A/*18S value obtained from the wild-type BY4741 strain. Data acquisition and processing were done by StepOne Software (v. 2.3; Applied Biosystems).

### Western blotting analysis

Cells were grown in YPD to the mid-log phase and an aliquot corresponding to 10^8^ cells was used to prepare whole-cell extract following a standard trichloroacetic acid-based protocol. Proteins were resolved by sodium dodecyl sulfate polyacrylamide gel (8%) electrophoresis and transferred to a nitrocellulose membrane. The antibodies used were anti-Gag (1:2500 dilution; raised in rabbits by Chang lab) and anti-Act1p (1:5000 dilution; Cat #A5441, Sigma; an internal control). Secondary antibodies were IRDye^®^ 800CW goat anti-rabbit (Cat # 926-32211, LiCOR, 1:10,000 dilution) and Alexa Fluor^TM^ 680 goat anti-mouse IgG (H+L) (Cat # A21058, Invitrogen, 1:10,000 dilution).

### Flow-cytometry-based competitive fitness assays

We measured the fitness of the three tester strains (L-A^-^, L-A^+^, or L-A^++^) by competing them individually against a reference strain expressing *TDH2*::GFP and *ADK1*::mCherry (from J.-Y. Leu, IMB, Academia Sinica) under various stress conditions. To ensure the measurements were taken at the exponential-growth stage, the tester and the reference strains were first separately pre-cultured at 30°C in synthetic complete (SC) medium overnight and then diluted and refreshed in media for 5 h to reach 0.1 OD_600_ (∼10^6^ cells). Equal number of tester and reference cells, each in ∼100 µl, were seeded into the stress-condition medium for a final volume of 5 ml and then grown to 0.2 OD_600_. The ratio of the two competitors was quantified at the initial and the final time points using a fluorescence-activated cell sorter (FACSCalibur, Becton Dickinson). GFP- and mCherry-expressing cells (10,000) in the resulting cell mixture were counted. The time-zero control was also counted in the same manner. Three independent replicates for each fitness measurement were performed. All stress experiments were performed in SC medium containing 2% glucose unless stated otherwise: (1) SC at 30°C; (2) 0.2 M NaCl/30°C (3) 0.4 M NaCl/30°C; (4) 0.8 M NaCl/30°C; (5) pH 3.8 (citric-acid buffered)/30°C; (6) pH 6.0 (disodium-phosphate buffered)/30°C; (7) SC/21°C; (8) SC/0.07% glucose/30°C; and (9) 3% glycerol/30°C.

### Transcriptomic and GO analysis

Independent colonies from each of the three L-A^-^, L-A^+^, and L-A^++^ isolates were cultured and subjected to two independent transcriptomics protocols. Total RNA was isolated using MasterPure^TM^ Yeast RNA Purification Kit (Lucigen) and treated with DNase I at 37°C for 30 m. In the first approach, done at the Genomics Core Facility, Institute of Molecular Biology, Academia Sinica, rRNA was first depleted prior to library construction and subsequent sequencing. In the second approach, done at Welgene Biotech Co., Ltd. (Taipei, Taiwan), the poly(A) RNA was specifically enriched without depleting rRNA for library construction.

Sequencing was done using Illumina NovaSeqX-PLUS. All sequencing data were analyzed by bioinformaticians in the Bioinformatic Core Facility, Institute of Molecular Biology, Academia Sinica Taipei, Taiwan. Sequencing data were aligned to the yeast reference genome yeast_R64 by STAR read mapper (versionL2.1.11b). Differential-expression analysis was done using HTseq/RSEM. Transcripts with expression fold change ≥2 and an FDR-corrected *P*-value ≤ 0.05 (false-discovery rate of 5%) were chosen for further analysis. Differentially expressed transcripts were subjected to GO analysis in term of biological processes using LAGO (https://go.princeton.edu/cgi-bin/LAGO). To identify genomic variations among nine L-A^-^, L-A^+^, and L-A^++^ isolates (three from each isolate), sequencing data from the rRNA-depletion protocol were analyzed by DeepVariant and SnpEff methods. The RNA-seq datasets generated and analyzed in this study have been deposited in the Gene Expression Omnibus (GEO) under accession number GSE295208.

## Acknowledgments

We thank C. Boone for the yeast libraries and J.-Y. Leu for advice. T.-H.C. was supported by Ministry of Science and Technology (MOST 110-2311-B-001-042) and by Academia Sinica Grand Challenge Seed Grant (AS-GCS-110-03) and Innovative Research Project (AS-IR-112-02-A).

## Author contributions

T.-H.C. conceived and designed the project; W.-Y.H., C.-S.Y., H.-I.L., and L.T. conducted the experiments; T.-H.C., W.-Y.H., and C.-S.Y. analyzed the data; W.-Y.H. and T.-H.C. wrote the manuscript.

